# Naturalistic acute pain states decoded from neural and facial dynamics

**DOI:** 10.1101/2024.05.10.593652

**Authors:** Yuhao Huang, Jay Gopal, Bina Kakusa, Alice H. Li, Weichen Huang, Jeffrey B. Wang, Amit Persad, Ashwin Ramayya, Josef Parvizi, Vivek P. Buch, Corey Keller

**Author notes:** Co-senior authors (Josef Parvizi, Vivek P. Buch, Corey Keller). <u>Corresponding Authors</u> Corey Keller Assistant Professor Psychiatry and Behavioral Sciences Stanford University Medical Center; Yuhao Huang Post-doctoral fellow Department of Neurosurgery Stanford University Medical Center. <u>Disclosures</u>. The authors have no relevant disclosures to this study. <u>Funding</u>. This study was funded in part by NIH R21 (1R21MH134172-01A1).

## Abstract

Pain is a complex experience that remains largely unexplored in naturalistic contexts, hindering our understanding of its neurobehavioral representation in ecologically valid settings. To address this, we employed a multimodal, data-driven approach integrating intracranial electroencephalography, pain self-reports, and facial expression quantification to characterize the neural and behavioral correlates of naturalistic acute pain in twelve epilepsy patients undergoing continuous monitoring with neural and audiovisual recordings. High self-reported pain states were associated with elevated blood pressure, increased pain medication use, and distinct facial muscle activations. Using machine learning, we successfully decoded individual participants’ high versus low self-reported pain states from distributed neural activity patterns (mean AUC = 0.70), involving mesolimbic regions, striatum, and temporoparietal cortex. High self-reported pain states exhibited increased low-frequency activity in temporoparietal areas and decreased high-frequency activity in mesolimbic regions (hippocampus, cingulate, and orbitofrontal cortex) compared to low pain states. This neural pain representation remained stable for hours and was modulated by pain onset and relief. Objective facial expression changes also classified self-reported pain states, with results concordant with electrophysiological predictions. Importantly, we identified transient periods of momentary pain as a distinct naturalistic acute pain measure, which could be reliably differentiated from affect-neutral periods using intracranial and facial features, albeit with neural and facial patterns distinct from self-reported pain. These findings reveal reliable neurobehavioral markers of naturalistic acute pain across contexts and timescales, underscoring the potential for developing personalized pain interventions in real-world settings.

## Introduction

Pain disorders represent a major burden of disease and severely reduced quality of life^1^. Developing a comprehensive understanding of pain processing and dysregulation is essential for informing diagnosis, monitoring, and treatment strategies. Pain is a multifaceted experience that involves the integration of distributed brain networks responsible for sensory, affective, and cognitive processing^2^. Much of our current knowledge about central pain processing stems from neuroimaging studies examining healthy individuals exposed to transient, experimentally induced painful stimuli^3–5^. These efforts identified neural markers of physical pain^3,6^, pain-related negative affect^4,7^, and pain-induced facial expressions^5^. While the sensory aspects of painful stimuli appear encoded in early sensory networks^3,4^, the generalized negative affect associated with pain is represented in a distributed set of regions, including the thalamus, orbitofrontal cortex (OFC), insula, and anterior cingulate cortex (ACC)^3,4^. However, the extent to which experimentally induced pain accurately represents the heterogeneous and persistent pain experienced in everyday life remains an important question^8^. For example, while tonic pain induced by capsaicin may better mimic natural pain compared to phasic thermal stimuli^9^, the extent to which experimental stimuli mirror real-life pain experiences remains uncertain. Therefore, studying pain within naturalistic and ecologically valid settings is important to elucidate the similarities and differences between experimental and naturalistic pain states, advancing our understanding of pain and facilitating the translation of this knowledge into potential therapeutic interventions.

Behaviorally, facial expressions serve as reliable indicators of pain across mammalian species^10,11^. These automatic changes in facial behavior likely stem from the complex integration of nociceptive, sensory, cognitive, and affective processes^12^. Moreover, pain-related facial expressions are thought to encode multidimensional pain experiences and may in part reflect activity within the spino-thalamo-cortical network^13^. While much research has focused on facial expressions in response to experimental pain, there is increasing recognition of the value in understanding how these dynamics unfold during naturalistic acute pain. This understanding is particularly valuable for aiding pain recognition in non-communicative individuals, and for characterizing facial expressions as a meaningful output channel of pain-related neural activity in everyday scenarios.

The inpatient epilepsy monitoring unit (EMU) provides a unique window into studying the neurobehavioral underpinnings of naturalistic acute pain states. Patients undergoing invasive monitoring for seizure localization often encounter varying degrees of acute pain during their stay^14^. Two forms of acute pain can be assessed: 1) patients’ self-reported assessments of their current or ongoing acute pain can be quantified using routine verbal rating scales^15^ and 2) episodes of momentary pain observable through behavioral cues such as changes in facial expressions and vocalizations^15^. Notably, intracranial electroencephalography (iEEG) recorded in the EMU offers exceptional spatiotemporal resolution, capturing direct neuronal activity over prolonged periods ranging from days to weeks^16^. Simultaneous video recording allows for the concurrent quantification of facial expressions alongside neural recordings on similar timescales. While iEEG has facilitated the identification of precise brain networks responsive to thermal pain stimuli^17^, and chronically implanted electrodes have demonstrated the involvement of the OFC and ACC in decoding self-reported chronic pain^18^, a knowledge gap persists regarding the neural and facial behavioral dynamics underlying variations in naturalistic acute pain states.

Employing a naturalistic, data-driven approach, we evaluated the two forms of acute pain in the EMU setting: 1) longitudinal acute pain as measured by pain self-reports and 2) transient episodes of momentary pain as identified by observational cues. First, we delineated the physiological and behavioral characteristics of self-reported pain states. High pain states were characterized by higher blood pressure, more frequent pain medication use and differential activation of facial muscles. Using spectro-spatial features derived from intracranial recordings, we then constructed a binary classifier capable of decoding self-reported pain states. Classifier interrogation revealed that pain decoding relied upon a distributed pattern of brain activity involving mesolimbic regions, thalamus, and temporoparietal cortices. Classifier output was temporally stable in between similar self-reported pain states and was sensitive to pain onset or analgesia of pain. Pain decoding based on objective facial behaviors performed worse but was positively correlated with electrophysiological decoding performance. We then extended these findings to transient episodes of momentary pain, demonstrating that momentary pain can be reliably decoded from periods of neutral affect using either neural or facial features. Notably, the neural and facial markers of momentary pain were distinct from those of self-reported pain states. Taken together, we find that distributed brain activity forms a reliable representation of naturalistic acute pain states and is closely mirrored by changes in facial dynamics.

## Methods

### Participants

The study included 12 human participants (5 females, ages 27-58; **Supplementary Table 1**) who underwent surgical implantation of intracranial electroencephalography (iEEG) recording electrodes for clinical seizure localization. All participants provided informed consent, monitored by the local Institutional Review Board, in accordance with the ethical standards of the Declaration of Helsinki. The decisions regarding electrode implantation, targets, and duration were made entirely based on clinical grounds, independent of this investigation. Participants were informed that their involvement in the study would not affect their clinical treatment, and they could withdraw at any time without compromising their care. While no statistical method was used to predetermine the sample size, the final sample size is comparable to contemporary iEEG literature ^19–21^.

### Pain self-report

Participants were routinely asked by the nursing staff what level of pain they are experiencing currently from a scale of 0 to 10, with 0 being no pain and 10 being the most intense pain ever felt. The median frequency of pain reporting for each participant was every two hours (Range: 1.5 – 2.5 hours; **Supplementary Table 2**). The pain score, painful body site, and the time of the pain report were documented. If more than one body site was painful, then additional pain scores were recorded for each body site. In this case, an average of all pain scores across the painful body sites was taken to obtain a mean pain score for that period.

### Participant Inclusion Criteria

We selected participants for the study who had (1) at least thirty measurements of self-reported pain (2) the median pain score was not zero, which would indicate a mostly pain-free monitoring period and precludes further analysis, (3) a pain score range of at least 50% of the total possible pain range during the monitoring period, thereby providing sufficient variation in self-reported pain scores, and (4) absence of large structural abnormalities (encephalomalacia, mass lesion, hematoma) as identified on MRI. Across participants, the mean range of pain scores was 74% of total possible pain range. (**Supplementary Table 2**).

### Pain medications

In addition to pain scores, nursing staff routinely asked participants if they would like to take as-needed pain medication. Pharmacological interventions administered included: acetaminophen, hydromorphone, gabapentin, ketorolac, lidocaine patch, acetaminophen-hydrocodone, oxycodone, acetaminophen-oxycodone, and tramadol. The names of medications, dosages, and administration times were documented.

Alternatively, non-pharmacological approaches to pain management could be utilized at the nurses’ discretion or participants’ requests, including repositioning, ice packs, heat packs, or verbal reassurance.

### Physiology

As part of routine care, heart rate, respiratory rate, and blood pressure were measured every 4 to 6 hours. Physiological measurements taken within 5 minutes of pain assessments were matched. Any physiological measurements falling outside this 5-minute window were excluded from the analysis.

### Electrode registration and anatomical parcellation

Electrode location in 3D space was obtained from post-implant CT co-registered with the subject’s pre-operative MRI^19,20^. Anatomic location of each contact was determined using FreeSurfer-based automated parcellation, as described previously^22^. Electrodes were further anatomically categorized into the following regions: parietal cortex, lateral and basal temporal cortex, hippocampus, amygdala, thalamus, lateral prefrontal cortex, orbitofrontal cortex, anterior cingulate cortex, striatum and the insular cortex. Electrodes located outside of the brain were excluded. Electrodes located in the occipital cortex were also excluded due to infrequent sampling. For electrode visualization, the FreeSurfer average brain was used with coordinates in standard Montreal Neurologic Institute (MNI) space.

### Data acquisition and signal preprocessing

Using stereoelectroencephalography (sEEG), neural recording from implanted depth electrodes (Adtech Medical; centre-to-centre contact spacing of 3 mm) were sampled at 1024 Hz (Nihon Koden). sEEG preprocessing and analysis were performed using FieldTrip^23^. Preprocessing consisted of notch filtering, re-referencing, and bandpass filtering. Specifically, a fourth order notch filter was first used to attenuate line noise (60, 120, and 180 Hz), followed by a laplacian re-referencing scheme to minimize far-field volume conduction^24^. Delta (1–4 Hz), theta (4–8 Hz), alpha (8–12 Hz), beta (15–25 Hz), gamma (25–70 Hz) and high-gamma (70–170 Hz) signals were obtained by using an 8th order zero-phase IIR Butterworth filter^25^. The filtered signals were subsequently downsampled to 100 Hz for further processing.

### Behavioral annotation of momentary pain episodes

Raw video footage was reviewed by two evaluators for episodes of momentary pain. All footage review was completed during periods of wakefulness. Momentary pain were considered present if any of the following observational cues were met: 1) facial grimace, 2) verbal statements reflecting pain such as “ouch”, “that really hurts”, 3) defensive behaviors such as bringing a hand to the site of pain and 4) pausing an ongoing activity due to pain^15^. No minimum duration was specified. Periods of ‘neutral’ affect were also labeled for comparison. Neutral periods were defined as any periods where there was absence of any affective facial expressions (happy, sad, cry), and absence of any of the above criteria for painful behaviors. For both momentary pain and neutral events, if the start and the end time of the painful or neutral periods were different between the two evaluators, then the average time was taken. If a particular event was only labeled by one evaluator, then this event was re-reviewed by the other evaluator to reach consensus. A minimum of five momentary pain events was required for each participant to be included for subsequent analysis. The mean number of momentary pain events identified was 20 with a mean duration of 18s (**Supplementary Table 3**). Only participants #5 and #10 did not exhibit the minimum number of momentary pain events, and they were excluded from this analysis.

### Acute pain state classifier construction, training, and evaluation

To investigate whether intracranial activity can predict self-reported pain states, we constructed a pain decoder based on spectro-spatial features. We primarily approached this as a binary classification problem by dividing pain reports into low versus high pain states on an individual participant level based on the median pain score.This was motivated by the observations that 1) people provide self-report ratings on a non-linear scale ^26^, 2) decoding of non-binary linear pain distributions performed much worse than binary pain classes in a cohort of chronic pain patients^18^ and 3) current clinical systems utilize a dichotomized signal threshold for closed loop stimulation paradigms (Medtronic Activa PC+S, Medtronic Percept). We constructed the spectro-spatial intracranial feature set by obtaining the log spectral power (root mean square) of the filtered signals for each channel and frequency band 5 minutes prior to the pain report. We evaluated signals prior to the report to avoid the confound of the neural signal being altered while participants reflected on their pain level. The 5-minute duration was chosen as a reasonable timeframe in which pain level is not expected to fluctuate dramatically. Due to the large number of potential spectro-spatial features present in the data (e.g., channels and powerband combinations), we employed Elastic-Net regression, a hybrid of ridge regression and lasso regularization favored in the presence of highly correlated variables, which is often the case with intracranial recordings^27^. To avoid data leakage while optimizing hyperparameters in the feature pruning process, we used a nested k-fold cross-validation (CV) scheme (**Figure 2B**). The number of folds, k, was either five or ten, ensuring that each fold contained at least five observations (pain measurements).In this scheme, data is split into a training set comprising k-1 fold of spectro-spatial features and pain state and a test set of the remaining fold. We used SMOTE (Synthetic Minority Over-sampling Technique) to upsample the class with fewer observations when present to account for class imbalance in the training set^28^. As a part of the inner *k*-fold CV scheme, we optimized the regularization strength of the Elastic-Net model. The optimal model based on the training set was then evaluated in the left-out test set. This process was repeated k times as a part of the outer k-fold CV scheme. Finally, a bootstrap was performed to allow for performance stability by repeating the process with random selection of cross validation indices each time for a total of 100 bootstraps. The null distribution is constructed by repeating the process with the pain label shuffled. Area under the curve (AUC) and accuracy were obtained as performance measures. As an ancillary analysis, we also trained multivariate linear models to predict continuous pain scores using the same k-fold CV scheme, with Elastic-Net models trained on continuous pain outcomes rather than binary outcomes. Pearson’s correlation value between actual and predicted pain scores was used as the performance measure.

### Neural classifier feature analysis

To evaluate the importance of iEEG spectro-spatial features, we analyzed the model coefficients obtained from the Elastic-Net trained models. Each feature’s coefficient can be either a non-zero number, indicating its inclusion in the model, or zero, indicating its exclusion. We sorted the mean feature coefficients from all 100 runs in descending order and estimated the change point in the cumulative summation curve based on mean and slope using the ’findchangepts.m’ function in Matlab. The cumulative set of features leading up to the change point were considered important. Furthermore, we assessed feature stability by determining the number of times each feature was selected across all 100 model runs. To ensure the robustness of the selected features, we compared the distribution of feature coefficients from the trained models with those from permuted models. Features whose coefficient distributions did not differ significantly between the trained and permuted models were discarded. This procedure was used previously to analyze intracranial neural features in the context of behavioral state classification analysis^21^. To investigate the relationship between spectro-spatial features and the underlying acute pain state, we first normalized the raw power values within each participant. We then computed the median power value for each feature across low and high pain states on an individual participant level. Finally, we pooled the median values across participants to derive the group average of feature values, stratified by either spectral band or anatomical region. This approach allowed us to identify the specific spectral and spatial characteristics of iEEG features that were most informative for decoding pain states.

### Neural classifier state timescale analysis

Next, we evaluated the stability of brain activity for making pain state classifications by calculating serial inferences using the index model trained on 5 minutes of spectro-spatial features immediately prior to pain self-reports. For each patient, we tested the stability of these neural features by using progressively distant non-overlapping 5-minute windows of data to calculate pain state probabilities without further model training. Rapidly changing neural activity since the index time should result in changes in the model output probabilities over time, whereas slowly changing neural activity should lead to relatively stable model output probabilities. To calculate the null serial probabilities, we trained the index model using shuffled pain state levels and subsequently used this model to make inferences. As the median time interval between pain self-reports ranged from 1.5 to 2.5 hours across participants, we limited the inferences to 3 hours after the time of pain self-report to prevent the model from encountering data on which it had already been trained. We stratified the pain state probabilities over time by the outcome class (low and high pain states) and categorized observations based on whether the subsequent pain measurement remained the same or changed. If a low pain state was followed by a high pain state, it indicated pain onset; conversely, if a high pain state was succeeded by a low pain state, it indicated pain relief or analgesia. Furthermore, we assessed the interventions used to alleviate pain within high pain states, including both pharmacologic and non-pharmacologic means.

### Non-linear pain state classification

To assess the performance of non-linear classification methods for pain states, we employed a random forest (RF) model, which utilizes multiple decision trees to converge on a classification output. RF models have been previously used successfully in decoding naturalistic affect^21^. For the RF classifiers, we specified the following hyperparameters based on a prior study^21^: 300 trees, the maximum number of samples per leaf between 1 and 20, and the number of features at each node between 1 and the total number of features minus 1. Similar to the Elastic-Net classification, we utilized a k-fold cross-validation scheme to evaluate the RF model’s performance. To determine feature importance, we used the out-of-bag error estimate derived from the RF models. This approach assesses the significance of individual spectro-spatial features by constructing each tree using bootstrap samples from the original dataset while reserving one-third of the data as a test set. Subsequently, the tree is constructed, and the omitted samples are classified, measuring the frequency of misclassification termed as the ’prediction error’. This process is a standard method for assessing feature importance in RF models^29^. To rank the features based on their importance, we performed change point identification on the prediction error for each feature, similar to the approach used for Elastic-Net features. We then compared each feature’s prediction error to its null distribution, which was obtained by shuffling the outcome labels across 100 runs. This process allowed us to identify the most informative spectro-spatial features for non-linear pain state classification using the RF model.

### Facial dynamic analysis

To investigate behavioral changes underlying naturalistic pain states, we evaluated facial dynamics, which have been extensively studied in the context of experimental and clinical pain^11^. We estimated moment-to-moment facial action units (AUs) from clinical videos recorded simultaneously with iEEG recordings. All participants in the epilepsy monitoring unit were routinely recorded with high-definition audio at 48 kHz and RGB video (640 x 480 pixels at 30 fps) 24 hours a day. The camera, located on the room ceiling, was manually centered on the participants’ faces by video technologists throughout their stay. This face-centered video capture is ideal for automated facial quantification approaches, which leverage deep learning models trained on large databases of mostly frontal-facing faces^30^. We partitioned the videos into non-overlapping 5-minute windows to mirror the parameters used for neural analysis. In each window, we computed estimates for common facial AUs using a multi-step process. First, we identified all faces in each video frame using a facial detection model (MTCNN). As family and staff were frequently present in the participant’s room, we isolated the participant’s face from the list of detected faces using a facial recognition model (DeepFace, https://github.com/serengil/deepface).

We then applied a facial AU model (OpenGraphAU^31^) to make inferences on a set of 27 facial AUs (AU1, AU2, AU4, AU5, AU6, AU7, AU9, AU10, AU11, AU12, AU13, AU14, AU15, AU16, AU17, AU18, AU19, AU20, AU22, AU23, AU24, AU25, AU26, AU27, AU32, AU38, AU39). This model provided the probability of each AU being present on the target face. From these frame-level features, we generated temporal-level features by computing the presence and average duration of each AU within a 5-minute time window. An AU was considered active if its inference probability was > 0.5 and lasted for at least 1 second, helping to eliminate spurious inferences on isolated frames. We used the Py-Feat toolbox to visualize the facial AUs that differed between low and high self-reported pain states.

Mirroring the classification approach used with electrophysiology, we trained an Elastic-Net regularized logistic regression model using facial features to classify between low and high self-reported pain states, as defined previously. We employed a k-fold cross-validation scheme to determine model performance. To evaluate the stability of facial activity in making pain state classifications, we calculated serial inferences based on the index model trained on 5 minutes of facial features immediately prior to pain self-reports. For each participant, we used progressively distant non-overlapping 5-minute windows of facial data to make serial inferences, representing facial expression activity across increasingly distant video timeframes in 5-minute increments after the pain report. To calculate the null serial probabilities, we trained the index model using shuffled pain reports and subsequently used this model to make inferences.

### Momentary pain classification

To classify between periods of momentary pain and neutral activity, we split the labeled time periods into one-second bins and trained three models for comparison. First, we trained an optimized model using an Elastic-Net regularized logistic regression, as defined previously, utilizing all spectro-spatial features. The outcome variable was the one-second bins, labeled as either momentary pain or neutral activity. We employed a conservative sequential 5-fold cross-validation scheme to account for the temporal dependency of behavioral timepoints occurring close together, which could artificially increase model performance. In this scheme, bins used for training and testing were grouped in time, ensuring that all one-second bins belonging to a labeled event (momentary pain or neutral) were not split between folds. To address the imbalance between the number of one-second bins for neutral events and momentary pain (**Supplementary Table 3**), we randomly down-sampled the neutral bins to match the number of momentary pain bins during model training for each participant. Second, we hypothesized that spectro-spatial features encoding self-reported pain states would also contain shared information representing periods of momentary pain. Consequently, we trained a logistic regression model using only the spectro-spatial features retained by the index binary pain self-report state classifier (trained on 5 minutes of data prior to the pain report). As the number of features was sparse due to pre-selection by the index classifier, we employed a logistic regression model without regularization. Third, we repeated the procedure using features retained by the index binary pain state classifier trained on shuffled pain reports. These features were not supervised by the self-reported pain states.

Each model condition underwent 100 bootstrap iterations to ensure robustness. Similarly, we performed classification of momentary pain and neutral periods based on facial dynamics. We constructed features as facial AU probabilities averaged within one-second time bins and employed an Elastic-Net regularized logistic regression with a sequential 5-fold cross-validation scheme, as described above.

### Comparison of neural and facial features-based pain state decoding

We performed analyses to evaluate the relationship between acute pain decoding based on spectro-spatial features and facial dynamics, as well as to determine whether facial features provided additional information for acute pain decoding. First, we assessed the correlation between the decoding performances of the two modalities by conducting Pearson’s correlation analysis on the AUC values obtained from the spectro-spatial and facial dynamics models across participants. Next, we investigated whether the integration of facial features with spectro-spatial features could improve acute pain decoding performance. To do this, we identified the spectro-spatial features retained by either the index binary pain self-report state classifier or the optimized momentary pain model. We then calculated the classifier performance using these features through cross-validation. Subsequently, we integrated the facial features into the neural feature set and evaluated the classifier performance using the combined set of neural and facial features, again employing cross-validation. By comparing the performance of the classifiers using spectro-spatial features alone with the performance of classifiers using the combined feature set, we evaluated whether facial features contributed additional, complementary information for acute pain decoding independent of neural features.

## Results

### Study participants and naturalistic pain states

We designed a naturalistic paradigm to study the neurobehavioral markers of acute pain states in an unconstrained manner, involving participants with implanted depth electrodes for seizure localization (**Figure 1A**). The study included regular pain self-reports, intracranial electrophysiology, and simultaneous audiovisual recordings. On average, we analyzed 6.3 days (SD = 2.1) of behavioral and electrophysiological data per participant, with channel coverage spanning cortico-thalamo-limbic networks (**Supplementary Table 1**). In addition to pain self-reports, we manually annotated periods of momentary pain based on video as another measure of naturalistic acute pain.

**Figure 1:**
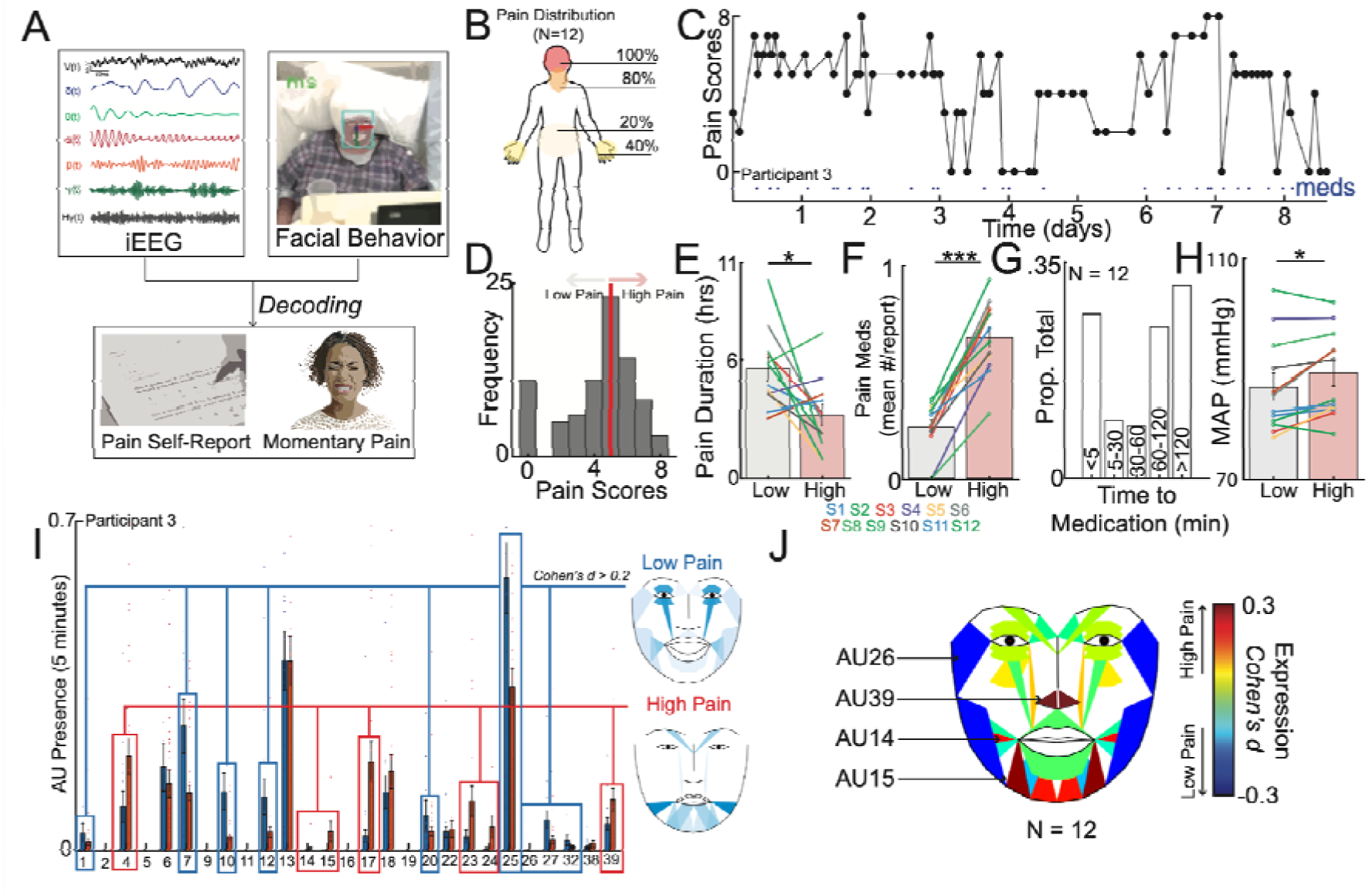
Naturalistic study design and inpatient tracking of acute pain states. **A)** The naturalistic paradigm comprises recording intermittent verbal pain self-report along with simultaneous intracranial electroencephalography (iEEG) and patient videos. Transient episodes of momentary pain are manually annotated based on video review. iEEG spectro-spatial features and quantitative facial dynamics are used to decode acute pain states. **B)** Group-level anatomical distribution of self-reported pain locations (N = 12). **C)** Variations in pain scores for an example participant over the course of nine days. Dots denote time when a pain medication was given. **D)** Histogram of all recorded pain scores for the same example participant. The median value was used to denote the threshold to define low versus high acute pain states. **E)** Amount of time spent in low versus high pain states based on consecutive pain reports that are of the same state. Participants overall spent less time in the high pain state compared to the low pain state (paired t-test; t(11): 2.6, P=0.02). **F)** More pain medications were given during the high pain state compared to the low pain state (paired t-test; t(11): 10.1, P<.001). **G)** Distribution of pain medication timing with relations to time of pain report. **H)** Higher mean arterial pressure (MAP) across participants was observed in the high pain state as compared to the low pain state (paired t-test: t(11): 2.7, P=0.02). **I)** Proportion of different AUs encountered during high versus low pain states for an example participant. Certain AUs denoting a positive affect are more expressed during low pain states whereas AUs associated with negative affect are more expressed during high pain states. **J)** Consensus AUs (*d* > 0.2) across participants which were differentially expressed between acute pain states. Colors represent the effect size between high versus low pain states. *Denotes P < 0.05, **P < 0.01 and *** P < 0.001.

**Figure 2:**
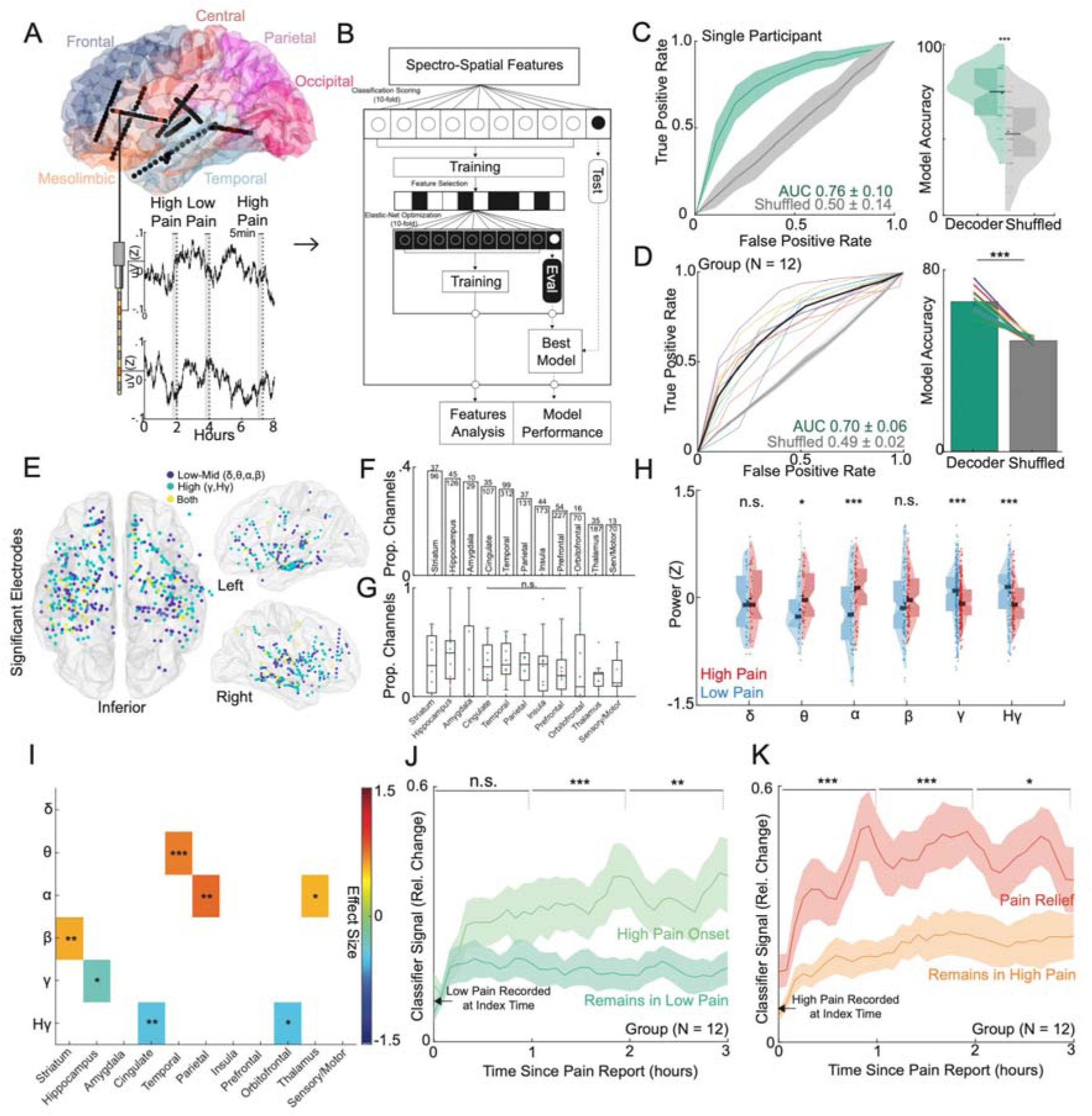
Intracranial neural activity sufficient for decoding of self-reported acute pain states are spatially distributed, temporally stable and modulated by pain onset or pain relief. **A)** Location of implanted depth electrodes in an example participant. Traces of raw voltage recording are shown for the two-colored electrodes over an 8-hour period, during which three self-reported pain scores were recorded. Five minutes prior to each pain score are used to construct spectro-spatial features. **B)** Spectro-spatial features are subsequently used to train an Elastic-Net regularized logistic regression model to classify between low versus high pain states. A nested cross-validation design is used to optimize hyperparameter selection and prevent overfitting. C) *Left Panel:* Mean receiver operating characteristics (ROC) curve for the same example participant for self-reported pain state classification across cross-validation folds and bootstraps. The grey curve represents the ROC curve when the outcome label is randomly shuffled. The shaded error bar represents the s.e.m across cross-validation folds and bootstraps. *Right Panel:* Accuracy of the model in prediction of pain states across folds and bootstraps. Accuracy of the model is significantly higher than the shuffled model (two-sample t-test; P<0.001). D) *Left Panel:* Group ROC curve for prediction of self-reported pain states across 12 participants. The shaded error bar represents the s.e.m across 12 participants. *Right Panel:* Accuracy of the model in prediction of pain states across 12 participants. Accuracy of the model is significantly higher than the shuffled model (paired t-test: t(11):10.2, P<0.001). **E)** Significant electrodes across participant classifiers are shown on a common brain in MNI coordinates. Color indicates whether a low to mid frequency (delta, alpha, theta, beta), high-frequency (gamma, high-gamma) or both type of spectral features were used at that location. **F)** Bar graph showing the sorted proportion of different anatomical regions recruited by classifiers across participants. **G)** Box plots showing proportion of different anatomical regions recruited stratified by individual participants. Each color dot represents a single participant. No significant difference was observed in the proportion of recruited electrodes across anatomical regions (Chi-square; χ2 = 7.2, P = 0.70). **H)** Normalized median distributions of low versus high pain state feature values as stratified by power bands. The median values from high pain states were significantly different from low pain states within theta (two-sample t-test; t(86): 2.41, P=0.02), alpha (t(138): 4.81, P<0.001), gamma (t(510): 3.25, P<0.001) and high gamma (t(234): 3.21, P<0.001) power bands. **I)** Heatmap displaying the power-region feature pairs that demonstrate a consistent effect size between high and low pain states (one-sample t-test; Single asterisk is p<0.05, double asterisk is p<0.01, and triple asterisk is p<0.001; FDR correction for multiple test comparisons). **J)** Percent change in the index classifier signal overtime when starting in a low pain state and subsequently stratified by if the next pain measurement remains in a low pain state or transitions to a high pain state (pain onset). Greater percent change in classifier signal is observed during pain onset. Shaded error bars represent s.e.m across participants. **K)** Percent change in the index classifier signal overtime when starting in a high pain state and subsequently stratified by if the next pain measurement remains in a high pain state or transitions to a low pain state (analgesia). Greater percent change in classifier signal is observed during analgesia. Shaded error bars represent s.e.m across participants. Single asterisk is p<0.05, double asterisk is p<0.01, and triple asterisk is p<0.001

To characterize acute pain states, we first analyzed regular pain self-reports obtained from participants. An average of 58 (SD = 21) pain self-reports were documented per participant, with a median frequency of every 2 hours (IQR = 0.3) (**Supplementary Table 2**). The most frequently reported pain locations were the head and jaw, followed by the hands and lower back (**Figure 1B**). In an example participant, visualizing all reported pain scores over time revealed fluctuations within and across days (**Figure 1C**; **Supplementary Figure 1** for other participants). We subsequently defined low and high acute pain states using the median pain score as the division threshold (**Figure 1D**). This approach was motivated by findings that most individuals do not reliably rate pain on a linear scale^26^, binary decoding of chronic pain performs better^18^, and a dichotomized signal threshold might be more clinically practical. Across participants, the median duration of pain states was 4.3 hours (IQR = 7.1) based on when pain scores transition from one state to another, with participants spending more time in low compared to high pain states (**Figure 1E**; paired t-test; t(11) = 2.6, P = 0.02). We also evaluated pain medication use surrounding the times of pain self-reports. More pain medications were administered within a 1-hour window centered on high pain states compared to low pain states (**Figure 1F**; paired t-test; t(11) = 10.1, P < 0.001). The timing of pain medications was heterogeneous, with some scheduled and others given as needed, resulting in a large distribution of medication timing relative to pain self-report times (**Figure 1G**). Participants in high pain states experienced higher blood pressure compared to low pain states (**Figure 1H**; paired t-test: t(11) = 2.7, P = 0.02), but no significant changes were observed for heart rate (**Supplementary Figure 2A**; one-sample t-test; t(11) = -1.50, P = 0.16) or respiratory rate (**Supplementary Figure 2B**; t(11) = - 1.93, P = 0.08).

Moreover, we quantified changes in facial behaviors using continuous video recordings to understand behavioral aspects underlying self-reported pain states. Facial behaviors were derived from frame-by-frame estimates of facial action units (AUs), which taxonomize human facial movements (see *Methods*). In an example participant, low pain states had higher expression of facial AUs denoting positive affect, whereas high pain states had higher expression of facial AUs denoting negative affect (**Figure 1I**). Across participants, several facial AUs were differentially expressed between acute pain states, with AU14, AU15, AU26, and AU39 exhibiting consistent effect sizes (**Figure 2J**; Cohen’s *d* > 0.2).

### Self-reported acute pain states decoded from distributed intracranial activity

To explore the neural mechanisms underlying self-reported acute pain states, we leveraged multi-site semi-chronic iEEG recordings (**Figure 2A**). Five-minute windows prior to pain self-report times were used to construct spectro-spatial features, which were subsequently employed to train an Elastic-Net classifier for decoding acute pain states (**Figure 2B**). We utilized a nested *k*-fold CV scheme to ensure that model performance was purely determined on unseen test data for each individual participant (**Figure 2B,C**, see *Methods*). At the group level, we achieved a mean classification AUC of 0.70±0.06 and a mean accuracy of 66%±5.5 for discriminating between low and high acute pain states (**Figure 2D**). We also evaluated the classification performance of a non-linear model (Random Forest, see Methods), as discrepancies in performance may suggest underlying non-linear data statistics. RF models performed slightly better, with a mean classification AUC of 0.72±0.06 and a mean accuracy of 69%±6.3 (**Supplementary Figure 3**). Moreover, we performed Elastic-Net based decoding of continuous pain scores (**Supplementary Figure 4**). Median Pearson’s R between actual and predicted pain scores was 0.29 (range: 0.01 to 0.70), with six out of twelve participants achieving significant decoding. In contrast, eleven participants exhibited significant binary decoding (**Supplementary Table 4**). Given the highly variable performance with continuous decoding, these results supported pain dichotomization, and subsequent analyses focused on results from the binary classifiers.

### Intracranial neural activity sufficient for decoding of self-reported acute pain states are spatially distributed, temporally stable and modulated by pain onset or pain relief

To elucidate the spectro-spatial contributions to the classification of self-reported acute pain states, we examined the spatial profiles of significant neural features (see *Methods*). We visualized the locations of each recruited electrode and its associated spectral content (**Figure 2E**), revealing a distributed spatial recruitment pattern with a mix of low-to-mid frequency bands (delta, theta, alpha, beta powers) and high-frequency bands (gamma, high-gamma). Fewer electrodes contained predictive spectral content spanning both low-to-mid and high frequencies for pain states. The anatomical distribution of significant electrodes was widespread, with the striatum, hippocampus, amygdala, cingulate cortex, and temporal cortex harboring the highest proportion of features predictive of pain states (**Figure 2F**). However, the proportion of significant features did not differ significantly across regions (**Figure 2G**; Chi-square test: χ2 = 7.2, P = 0.7), highlighting the distributed nature of acute pain markers. Irrespective of feature normalization or classifier type, the gamma band was the most common predictive frequency band (**Supplementary Figure 5**).

Across all significant spectro-spatial features, high pain states exhibited a consistent increase in theta power (linear regression coefficient adjusted for participant; t(85) = 2.44, P = 0.02) and alpha power (t(137) = 4.81, P < 0.001) compared to low pain states, accompanied by a decrease in gamma (t(509) = 3.26, P < 0.001) and high-gamma power (Figure 2H; t(233) = 3.21, P < 0.001). When stratified by anatomical region, the aggregate increase in theta and alpha power was driven by temporal, parietal, and thalamic regions, while the decrease in gamma and high-gamma power originated from mesolimbic regions (Figure 2I; linear regression coefficient adjusted for participant: * < 0.05, ** < 0.01, *** < 0.001, FDR-correction for multiple test comparisons). Note that while other regions did not exhibit consistent directional changes in power, they were still important in predicting acute pain states. To assess the consistency of these neural feature patterns, we performed feature analysis using random forest classifiers (see *Methods*). The random forest classifier relied on distributed neural activity, with a similar pattern of increased theta (t(173) = 2.1, P = 0.04) and alpha (t(241) = 5.8, P < 0.001) power, as well as decreased gamma (t(427) = 3.5, P < 0.001) and high-gamma (t(231) = 4.4, P < 0.001) power (**Supplementary Figure 6A-B**). However, we also noted a decrease in delta power (t(221) = 5.6, P < 0.001) and an increase in beta power (t(271) = 6.2, P < 0.001). Anatomically, similar decreases in high-frequency activity were observed in the hippocampus and anterior cingulate cortex, while increases in theta and alpha power were seen in the temporal and parietal cortices (**Supplementary Figure 6C**; linear regression coefficient adjusted for participant: * < 0.05, ** < 0.01, *** < 0.001, FDR-correction for multiple test comparisons). In summary, neural activity underlying self-reported acute pain decoding is distributed, exhibiting regionally specific spectral profiles.

We next evaluated the temporal stability of acute pain neural encoding between pain self-reports. We hypothesized that if self-reported pain states lasted on the timescale of hours, the output of the pain state classifier should remain relatively stable when using unseen spectro-spatial features over a similar time period. Additionally, if participants underwent a self-reported pain state transition, the classifier should be sensitive to that change. Starting with observations recorded as low pain, across participants, when the next pain measurement remained in a low pain state, the relative change in classifier signal was fairly stable (mean % change across time: 0.19; **Figure 2J**), indicating that neural features encoding pain states are stable in the absence of state change. In contrast, when the next pain measurement transitioned to a high pain state, we observed a larger change in classifier signal over time (mean % change across time: 0.31), which was significantly different compared to when the pain measurement remained stable (**Figure 2J**; paired t-test: t(11) = 2.64, P = 0.02). Similar findings were observed when starting with high pain state observations (**Figure 2K**). The classifier signal remained stable over time when the next measurement stayed in a high pain state (mean % change across time: 0.25). However, when pain relief was experienced (i.e., the next measurement transitioned to a low pain state), a significantly larger change in classifier signal was observed (mean % change across time: 0.39; t(11) = 3.0, P = 0.01). Since pain medications were often administered during high pain states, we evaluated the effect of medications given at the time of a high pain self-report (**Supplementary Figure 7**). We found that medications alone did not change the classifier signal. For instance, medications given without a self-reported change in pain state did not result in a classifier signal change (t(11) = 1.82, P = 0.1). Medications given with associated pain relief produced a signal change similar to that of pain relief achieved through non-pharmacologic means, such as rest, elevation, verbal reassurance, or physical comfort (t(11) = 1.1, P = 0.3). Notably, pain scores were rated on average 1 point higher across participants in high pain states when pain medication was given compared to high pain states in which non-pharmacologic interventions were used (**Supplementary Figure 8**). These results indicate that neural features underlying self-reported pain states change slowly, are tuned to state transitions, and are not directly altered by pain medications.

### Facial behavioral underlying self-reported acute pain states

Mirroring the analytical approach for neural-based acute pain decoding, we trained a classifier using five minutes of video-based facial features immediately prior to pain self-report (Figure 3A). Using facial features alone, we achieved a group AUC of 0.62 ± 0.09 in classifying self-reported pain states (**Figure 3B**). Notably, model performance based on facial dynamics was concordant with neural-based decoding at the group level; participants who demonstrated high AUC using neural activity to predict self-reported pain states also had high AUC using facial data to predict self-reported pain states (**Figure 3C**; Pearson’s R = 0.70, P = 0.01). We also evaluated the temporal stability of facial features in encoding pain states (**Figure 3D**). Starting with high pain state observations, the facial feature-based classifier demonstrated a greater signal change when the next measurement indicated pain relief compared to remaining in a high pain state (paired t-test; * < 0.05, ** < 0.01, *** < 0.001). However, the facial feature-based classifier was not sensitive enough to detect differences between remaining in a low pain state and pain onset (**Supplementary Figure 9**). Given the observed correlation between facial and neural-based decoding, which suggests a relationship between neural features and facial expression changes, we investigated whether facial features contributed unique predictive value or redundant information. We compared the classifier AUC using neural features alone to the AUC when neural and facial features were combined. The inclusion of facial features did not improve the decoding of self-reported pain states (**Supplementary Figure 10**; paired t-test: t(11) = 0; P = 0.99). These findings demonstrate that facial dynamics alone can be used to classify acute pain states, albeit with lower performance compared to neural-based decoding. The concordance between facial and neural-based decoding at the group level suggests a shared underlying representation of pain states. However, the lack of improvement in decoding performance when combining facial and neural features indicates that facial features may not provide additional predictive value beyond what is already captured by neural activity.

**Figure 3:**
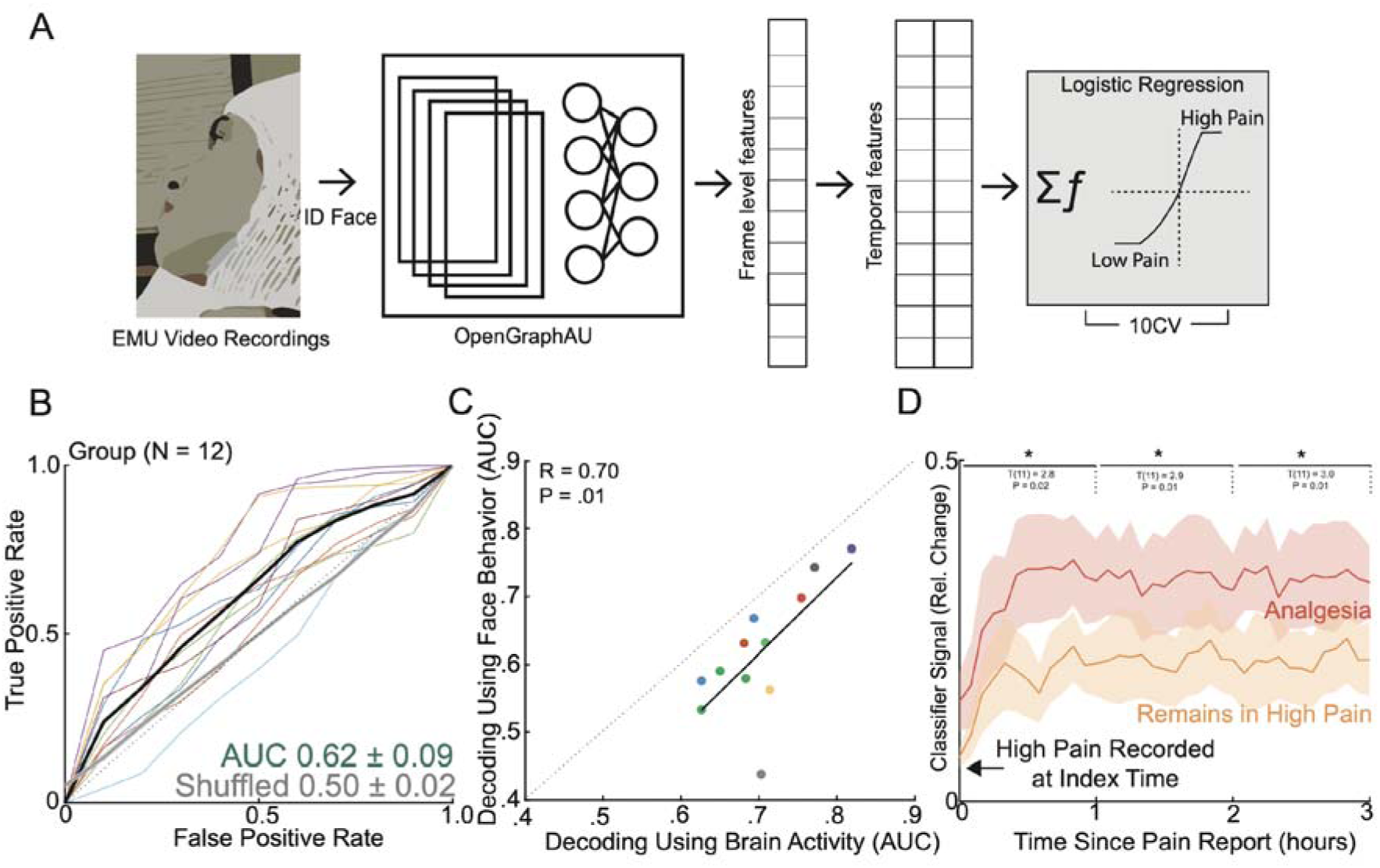
Facial dynamics underlying acute pain states. **A)** To evaluate aspects of behavior during high versus low self-reported pain states, we performed automatic quantification of facial muscle activation on a per-video frame basis. To do this, we devised a custom video processing pipeline where faces were first extracted from a frame, then the participant of interest was isolated from staff and family members using facial recognition, and finally the isolated face embeddings were fed into a pre-trained deep learning model to facial action unit (AU). Frame level AU outputs were collapsed across time to generate temporal statistics, which were subsequently used to decode self-reported pain states in a nested cross-validation scheme. **B)** Group ROC curves for decoding pain states using facial dynamics quantified during the five-minute window prior to pain report, which is the same time window used for electrophysiological pain decoding. Lines represent participants while the shaded gray error bar represents the decoding performance when the outcome label was shuffled. **C)** Performance of decoding based on facial dynamics is directly correlated with decoding based on brain activity (Pearson’s R: 0.70, P=0.01). Given all the points are on the right of the diagonal, brain decoding outperforms facial behavioral decoding for all participants. **D)** Percent change in the index classifier signal overtime when starting in a high pain state and subsequently stratified by if the next pain measurement remains in a high pain state or transitions to a low pain state (analgesia). Greater percent change in classifier signal is observed during analgesia. Shaded error bars represent s.e.m across participants. Single asterisk is p<0.05, double asterisk is p<0.01, and triple asterisk is p<0.001

### Neurobehavioral markers of momentary pain

As another measure of naturalistic acute pain, we manually annotated transient episodes of momentary pain based on observational cues (**Figure 4A**; see *Methods*). Across participants, we identified a total of 377 behavioral events (N_momentary pain_ = 203; N_neutral_ = 174), which occurred on average 34 minutes from the closest pain self-report (**Supplementary Figure 11A**). Labeled behavioral events were found to be equally close to either low or high self-reported pain states (**Supplementary Figure 11B**; χ² = 0.084, P = 0.77). Analysis of facial expression changes during momentary pain revealed several consensus changes across participants, including AU5, AU6/7, AU9/10, AU11/12, and AU25/26 (**Figure 4B**; Cohen’s *d > 0.2*). Notably, these action units have been previously identified as pain-related facial markers in studies of elicited pain^11^. In contrast, the facial action units associated with different self-reported pain states were AU14, AU15, AU26, and AU39 (**Figure 1J**), which are more affect-related. This difference in facial expression patterns suggests that momentary pain episodes may be more similar to conventionally elicited pain in experimental conditions, while self-reported pain states reflect a more complex and integrative experience of pain.

**Figure 4:**
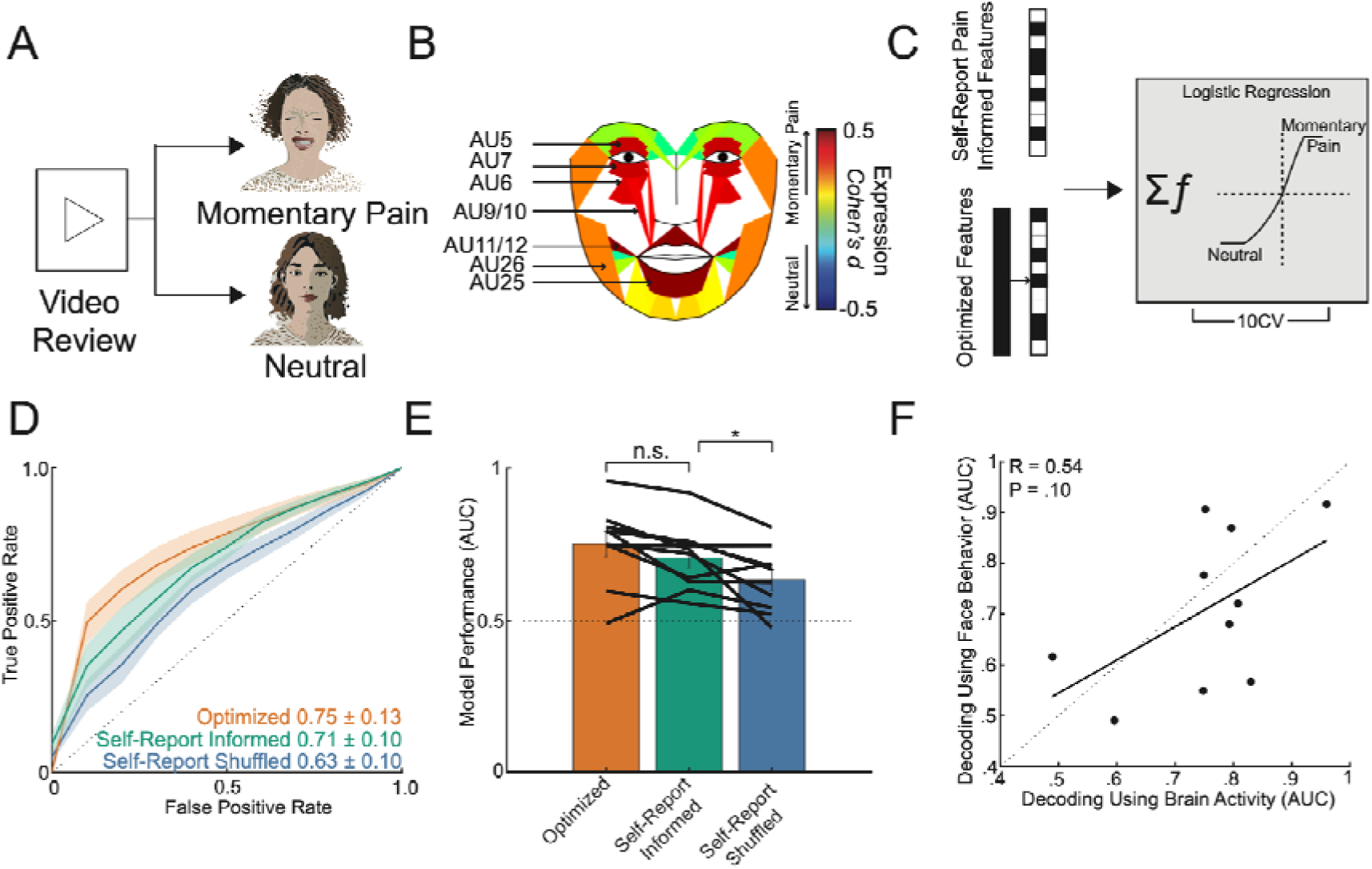
Transient episodes of momentary pain can be decoded using neural and facial activity. **A)** Manual video reviews by two evaluators were performed to identify periods of momentary pain and periods of neutral affect. **B)** Consensus AUs (*d* > 0.2) across participants which were differentially expressed between periods of momentary pain and periods of neutral affect. Majority of these AUs have been implicated in pain expression previously. Colors represent the effect size between momentary pain and neutral periods. **C)** Classifiers were trained using the full spectro-spatial feature set (optimized model) and using solely features previously selected by the index pain self-report state classifier (pain self-report informed model). **D)** ROC curves for prediction of momentary pain from neutral affect across participants, stratified by the optimized model, the pain self-report informed model, and the self-reported pain shuffled model (spectro-spatial features selected from pain self-report state classifier trained on shuffled pain labels). The shaded error bar represents the s.e.m across 10 participants. **E)** Bar plot comparison of the three trained models. There was no significant difference between the optimized and the pain self-report informed model performance (paired t-test; t(9): 2.0, P=0.07). The pain self-report informed model performed better than using features not supervised to the self-report pain states (paired t-test; t(9): 2.7, P=0.03) **F)** Comparison of momentary pain decoding based on facial dynamics and neural features.

To investigate the similarities and differences in neural features underlying momentary pain compared to self-reported pain, we trained three different classifier models (**Figure 4C**): 1) an optimized model using all available spectro-spatial features, 2) a self-report pain informed model using spectro-spatial features previously retained in the index self-report pain state classifier, and 3) a self-report pain shuffled model using spectro-spatial features previously retained in the index self-report pain state classifier but with shuffled pain labels. The self-report pain shuffled model represents a random set of neural features that are approximately equal in length to the features in the self-reported informed model and serves as a control. The optimized model achieved a group mean AUC of 0.75 ± 0.13 for classifying momentary pain periods from neutral affect (**Figure 4D**), while the self-report pain informed model achieved a group mean AUC of 0.71 ± 0.10. In contrast, the self-reported pain shuffled model achieved a lower group mean AUC of 0.63 ± 0.10. Although the optimized model did not significantly outperform the self-report pain informed model (**Figure 4E**; paired t-test: t(9) = -2.0, P = 0.08), the self-reported pain shuffled model performed significantly worse than the informed model (t(9) = - 2.7, P = 0.03). This finding indicates that neural features supervised to pain self-reports are more effective in classifying momentary pain than randomly selected neural features. To identify the spectral markers of momentary pain, we evaluated significant features in the optimized model. On an aggregate level, momentary pain states exhibited a decrease in theta (linear regression coefficient adjusted for participant; t(125) = 7.7, P < 0.001), alpha (t(93) = 4.6, P < 0.001), beta (t(107) = 5.8, P < 0.001), gamma (t(478) = 3.8, P < 0.001), and high-gamma (t(244) = 2.8, P = 0.01) power (**Supplementary Figure 12A**). Regionally, the decrease in gamma and high-gamma power was driven by hippocampal and prefrontal activity, whereas the decrease in theta, alpha, and beta power was observed in the cingulate, parietal, temporal, insular, prefrontal, and sensory/motor cortices (**Supplementary Figure 12B**). As an outlier to this pattern, the insular cortex demonstrated an opposite pattern, with increased gamma and high-gamma activity during momentary pain. We also assessed the decoding of momentary pain using facial dynamics (mean group AUC: 0.71±0.15) and compared it with neural-based decoding employing the optimized models. While there was a moderate correlation between the two decoding methods, it was not statistically significant (**Figure 4F**; Pearson’s R = 0.54, P = 0.10). Furthermore, incorporating facial features alongside neural features did not result in a significant improvement in model performance across participants (**Supplementary Figure 13**; paired t-test: t(9) = 1.2; P = 0.27). These findings demonstrate that momentary pain states can be reliably decoded, with some of the informative neural features overlapping with those involved in representing self-reported pain. However, the spectral markers of momentary pain differ from those of self-reported pain. This difference in spectral patterns suggests that momentary pain and self-reported pain may engage distinct neural processes, reflecting the multifaceted nature of pain experience in naturalistic settings. Finally, the lack of significant improvement in decoding performance when combining neural and facial features suggests that these modalities may capture overlapping information about momentary pain states.

## Discussion

The neural and behavioral correlates underlying everyday pain experiences are of significant clinical interest due to the substantial burden of prevalent pain disorders. While previous studies have provided insight in experimental contexts, there remains a paucity of studies evaluating naturalistic pain experiences in unconstrained real-world settings. Here, we examined the neural and facial correlates underlying acute pain using a purely naturalistic paradigm. This approach enabled us to longitudinally study the ecologically valid pain experience in a manner that would be ethically challenging to replicate experimentally. Our findings demonstrate that self-reported acute pain states over time can be accurately decoded using intracranial brain activity across participants. Neural features recruited for pain decoding were distributed across cortico-thalamo-limbic regions, with a bias towards high-frequency neural activity. Notably, high pain states were associated with increased low-frequency (theta, alpha power) activity in temporal and parietal areas alongside reduced gamma and high-gamma activity in mesolimbic regions including the hippocampus, cingulate, and orbitofrontal cortex. Furthermore, the self-reported pain classifier output was temporally stable and sensitive to changes in pain state.

Complementary facial analysis revealed discernible differences between self-reported acute pain states, concordant with neural decoding performance across participants. We also identified transient periods of momentary pain as a distinct naturalistic acute pain measure. Similar to self-reported pain, momentary pain could be reliably decoded from neutral states using intracranial and facial features, though with distinct neural spectral patterns compared to self-reported pain. These findings highlight robust neurobehavioral markers of naturalistic pain states across contexts and timescales, motivating further research into diagnostics and interventions of pain in unconstrained settings.

The distributed pattern of brain activity contributing to naturalistic pain decoding aligns with prior neuroimaging studies highlighting the involvement of various regions in pain processing^3–5,7^. These include areas specific to somatic sensations including the thalamus, posterior insula, and somatosensory cortex^32,33^, as well as regions relevant to general affect and salience such as the dorsolateral and ventromedial prefrontal cortices, parietal cortex, anterior insula, cingulate, and amygdalo-hippocampal complex^3,4^. Notably, our findings also revealed engagement of the medial and lateral temporal regions, consistent with a recent intracranial study where temporal gyri activity predicted thermal pain onset^17^. The temporal cortex has been proposed to encode memories associated with painful experiences, suggesting its involvement in the experiential aspects of pain processing ^34,35^. Additionally, striatal regions like the caudate nucleus and putamen have been implicated in modulatory pain systems^36^. Furthermore, various behavioral factors may contribute to the distributed neural representations of pain^37–39^. For instance, sympathetic arousal indexed by blood pressure elevations during self-reported pain^40^, as well as unmeasured states like changes in mood, verbal output, or motor movement which often co-occur with pain^41^. Given these potential behavioral co-occurrences, we interpret our intracranial findings as biomarkers of naturalistic pain experiences rather than specific neural signatures of nociceptive processing. The distributed patterns likely reflect the sensory, affective, and cognitive-evaluative components inherent to acute pain states.

The spectral patterns associated with pain states remain an active area of investigation. Scalp EEG studies have suggested that alpha and beta oscillations may encode the intensity of noxious stimuli, while gamma oscillations in the prefrontal region could represent prolonged experimental pain experiences ^42–45^. However, intracranial studies examining spectral power in relation to pain are scarce. Caston et al. found predominantly increased high-frequency power in several regions, notably the lateral temporal cortex and hippocampus, in response to thermal pain onset^17^. Conversely, Shirvalkar et al. demonstrated that chronic pain states were largely inversely related to oscillatory activity in the anterior cingulate (ACC) and orbitofrontal cortices (OFC), with a mixed relationship for experimental pain^18^. In our study, we observed differential spectral patterns associated with different measures of naturalistic acute pain. High self-reported pain states were linked to increased low-to-mid frequency power in the striatum, thalamus, and temporo-parietal regions, alongside decreased high-frequency power in the hippocampus, cingulate, and OFC. In contrast, momentary pain was associated with broadly decreased power across frequencies, particularly high-frequencies in the hippocampus and prefrontal regions, and low-to-mid frequencies in other areas.

These spectral differences may reflect the nature of the pain experience - self-reported pain states may encompass a broader range of pain-related sensations, emotions and cognitive processes represent temporally stable, integrated processing of pain-related sensations, emotions and cognitive processes, whereas momentary pain captures immediate, highly salient pain events akin to experimental stimuli. Indeed, facial analyses indicated differential muscle activations during high self-reported pain versus momentary pain, suggesting distinct experiential qualities. Together, these findings highlight the complex spectral dynamics underlying naturalistic acute pain experiences.

The naturalistic design of our study provided a unique opportunity to investigate the temporal dynamics of neurobehavioral pain markers. We found that between stable pain self-reports, both the neural and facial markers of pain remained relatively constant, without exhibiting rapid fluctuations. In contrast, when the subsequent pain self-report indicated either pain onset or pain relief, these pain-related markers demonstrated gradual changes, with neural markers showing greater sensitivity compared to facial markers. These results suggest that self-reported pain states over time operate on longer timescales, likely reflecting integrated processing of the current aversive state rather than rapid encoding of sensory/nociceptive information. The temporal stability of pain markers between self-reports, coupled with their gradual modulation preceding state changes, highlights the sustained neural and behavioral representations of the subjective experience of acute pain over time in naturalistic settings.

Behaviorally, we used quantitative facial mapping to assess the facial correlates of self-reported pain states as well as momentary pain. While many prior studies have evaluated facial action unit (AU) differences in relation to pain in controlled settings^46,47^, we found that facial expression changes underlying self-reported pain states had little overlap with conventionally reported pain-related AUs. Instead, these AUs (AU14, AU15, AU26) may reflect more general changes in affect ^48^. In contrast, facial expressions associated with momentary pain exhibited many of the canonical pain-related AUs (AU5, AU6/7, AU9/10, AU11/12, AU25/26), suggesting immediate pain onset. Notably, we observed concordance between the facial and neural decoding of acute pain states, implying that some of the informative neural activity is related to the outward manifestation of pain. This is further supported by our finding that facial features did not independently contribute substantial performance gains, indicating that the neural features likely explained much of the variance in facial behavior. Prior literature has demonstrated correlations between neural activity and facial expressions during pain^12,13^. Additionally, the activity of regions such as the cingulate and amygdala is known to influence the regulation of emotional expressions^49^. These results highlight the tight coupling between the neural and facial representations of naturalistic acute pain experiences.

### Limitations and Future Directions

The recording locations in our study were solely determined by clinical requirements for epilepsy localization, resulting in variable electrode coverage across participants. This variability poses a challenge for developing generalizable models that can predict pain states across individuals, as electrodes are rarely positioned identically. An intriguing future direction would be to explore whether models trained on standardized scalp EEG data or large datasets of intracranial recordings with similar brain coverage could effectively predict pain states in diverse individuals.

Furthermore, our naturalistic study design precludes direct causal inferences, raising the possibility that the observed correlations between intracranial activity and acute pain states may be influenced by unmeasured variables such as sympathetic arousal, altered verbal output, or motor activity, as discussed previously. To address this limitation, future studies could employ direct electrical brain stimulation to causally manipulate specific regions across varying timescales while closely monitoring pain experiences. Such interventional approaches, combined with neural recordings, could elucidate the causal mechanisms underlying the neurobehavioral markers of naturalistic acute pain identified in our study. Additionally, incorporating multimodal physiology data such as skin conductance and pupillary tracking could provide a more comprehensive characterization of the multidimensional pain experience.

